# Cysteine is a limiting factor for glioma proliferation and survival

**DOI:** 10.1101/2021.01.08.425930

**Authors:** Victor Ruiz-Rodado, Tyrone Dowdy, Adrian Lita, Tamalee Kramp, Meili Zhang, Jinkyu Jung, Ana Dios-Esponera, Christel C. Herold-Mende, Kevin Camphausen, Mark R. Gilbert, Mioara Larion

**Affiliations:** Neuro-Oncology Branch, Center for Cancer Research, National Cancer Institute, National Institutes of Health, Bethesda, US; Radiation Oncology Branch, Center for Cancer Research, National Institutes of Health, Bethesda, US; Fred Hutchinson Cancer Research Center, Seattle, US; Division of Neurosurgical Research, Department of Neurosurgery, University Hospital Heidelberg, Heidelberg, Germany

## Abstract

Nutritional intervention is becoming more prevalent as adjuvant therapy for many cancers in view of the tumor dependence on external sources for some nutrients. We report the dependence of glioma cells on exogenous cysteine/cystine, despite this amino acid being nonessential. ^13^C-tracing and the analysis of cystathionine synthase and cystathioninase levels revealed the metabolic landscape attributable to cysteine deprivation, and the disconnection between the methionine cycle and the transsulfuration pathway. Therefore, we explored the nutritional deprivation in a mouse model of glioma. Animals subjected to a cysteine/cystine-free diet survived longer, with concomitant reductions in glutathione and cysteine plasma levels. At the end point, however, tumors displayed the ability to synthesize glutathione, although higher levels of oxidative stress were detected. We observed a compensation from the nutritional intervention revealed as the recovery of cysteine-related metabolites in plasma. Our study highlights a time window where cysteine deprivation can be exploited for additional therapeutic strategies.

## Introduction

The metabolism of tumor cells depends on many factors, notably, nutrient availability from the microenvironment and oncogenic mutations. Others have studied whether restricting the nutrient availability from the microenvironment can reduce the distinctive high proliferation rate of Myc-driven tumors, which require glutamine (1-3), or melanoma cells, which require leucine (4) or serine (5). Although such dietary interventions have been explored recently, with the goal to “starve” cancer cells (6, 7), their nutrient requirements and the mechanisms underlying their dependence on specific metabolites obtained from the environment are not fully understood. Gliomas are brain tumors that can have an aggressive phenotype and currently have no curative treatment. The potential of incorporating a dietary plan into the treatment of this disease is highly desirable by patients. In clinical studies of gliomas, one of the most popular diets as adjuvant therapy is the ketogenic diet (8), with over 12 clinical trials currently exploring that type of intervention (NCT03451799, NCT03328858, NCT03160599, NCT02694094). However, no other interventional diets are being explored for this disease, in part due to a lack of preclinical evidence. Gliomas harboring IDH1 mutations have been reported to have a decreased ability to compensate for reactive oxygen species (ROS) (9), due to reduced IDH1-wildtype activity and concomitant low nicotinamide adenine dinucleotide phosphate (NADPH) (10). Recent reports noted that upregulation of antioxidant pathways compensates for the increased ROS levels found in IDH1-mutant tumors (11). By tilting the balance towards increased oxidative stress (12), potentially via interventional diets, these findings provide a new framework to investigate alternative therapeutic strategies for gliomas.

Diets that restrict specific amino acids, particularly essential amino acids, have been explored in other cancers (6). However, cysteine is a nonessential amino acid, as it can be synthetized from methionine through the transsulfuration (TS) pathway. The most abundant form of cysteine is cystine, which has plasma levels 10 times higher than cysteine (13). Once cystine enters the cell it is reduced to cysteine, which then can be condensed with glutamate to form glutamylcysteine (Glu-Cys), a reaction that is the rate-limiting step for glutathione (GSH) synthesis, which is a major ROS scavenger. Alternatively, methionine can be converted into homocysteine through its sequential transformation into the metabolic intermediates S-adenosylmethionine (SAM) and S-adenosylhomocysteine (SAH). Subsequently, homocysteine can either be diverted into cysteine synthesis through the TS pathway or be remethylated to yield methionine and generate tetrahydrofolate. Hepatic tissue synthetizes almost half of the total pool of GSH from methionine-derived cysteine (14), although the importance of the TS pathway is not limited to the liver. Indeed, experiments conducted in mice revealed 29% less GSH in brain presenting homozygous cystathionine β-synthase gene disruption (15). However, it is not clear when this pathway plays a main role in supplying cysteine instead of being taken up from the environment (16).

Herein, we describe the metabolic landscape resulting from cysteine/cystine-deprivation, including metabolic alterations beyond cysteine metabolism. We then propose a new method to hypersensitize glioma cells to oxidative stress without using any drug treatment. We translated our approach to an *in vivo* model of IDH1-mutant glioma and found a transitory disruption of cysteine metabolism at the systemic level, which is compensated over time. Nevertheless, it provides a survival benefit in our animal model. This study highlights how we can use dietary interventions to create metabolic vulnerabilities that can be exploited to design more efficient therapy.

## Results

### Cysteine and cystine deprivation halt the growth and reduce the viability of gliomas by downregulating protein translation and glutathione synthesis

Three glioma cell lines (BT142, TS603, and NCH1681) were cultured in DMEM:F12, which either contained or lacked both cysteine and cystine. The absence of those amino acids had an antiproliferative effect (Figure 1A), which was more intense in the BT142 cell line, limiting its growth to 50% over the course of 96 hours. Moreover, cells were unable to form neurospheres (Figures 1B and S1A), and their viability was reduced by approximately 30% in 96 hours (Figure 1C). At 72 hours, all cell lines displayed a significant decrease (*p* < 0.001 for all cell lines) in viability, and TS603 even had a significant decrease in viability at 48 hours (*p* = 0.038). To examine the key factors involved in both the antiproliferative effect and the reduction in viability resulting from cysteine/cystine-deprivation, we conducted rescue experiments by treating the cells with downstream metabolites of cysteine or methionine, such as SAM, cystathionine, homocysteine, taurine, or GSH (Figure 1D). We also employed an inhibitor of the IDH1-mutant enzyme (AGI5198) because these cell lines harbor this mutation. This agent did not alter the deleterious effect of cysteine/cystine-deprivation (Figure 1E); indeed, IDH1-wildtype glioma cell lines were also affected by this treatment (Figure S1B). The experiments revealed that only homocysteine and cystathionine could restore cell growth; thus, we explored the link between their supplementation and an increase in cysteine availability by checking the levels of GSH and the expression of glutamate-cysteine ligase modifier subunit (GCLM) (Figure S1C), which acts as a sensor of cysteine levels (17) in GSH synthesis. Both GSH (Figure 1F) and the expression of GCLM were reduced under cysteine/cystine-deprivation, and their levels recovered after adding homocysteine and cystathionine, revealing that both metabolites are cysteine donors. Amino acid deprivation has been shown to inhibit protein synthesis in cancer (18), and because cysteine is also a proteinogenic amino acid, we evaluated the capacity of the cells subjected to cysteine/cystine-deprived conditions to synthesize protein. We examined the levels of phosphorylated eIF2α (p-eIF2α) and GCN2 (p-GCN2), two proteins involved in the stress response to amino acid starvation, and uncharged t-RNAs (19, 20) in addition to treating glioma cells with puromycin and then analyze the incorporation to nascent proteins, as previously described (21). p-eIF2α and p-CGN2 levels were both higher in the samples grown in the absence of cysteine/cystine (Figure 1G). Furthermore, after 72 hours, levels of puromycin-positive labeled proteins were higher for glioma cell lines grown in full medium than for cells deprived of cysteine/cystine (Figures 1H and S1E), indicating that deprivation reduces the protein synthesis activity. Moreover, both homocysteine and cystathionine recovered cell viability together with GSH and Trolox, an antioxidant analogue of vitamin E, suggesting that the loss in viability was linked to the decreased ability of these cells to buffer the naturally occurring ROS (Figures 1I and S1D) rather than to the availability of cysteine/cystine. However, the ability of GSH to provide full protection against ROS was constrained by its poor cellular permeability (22), since the intracellular levels of GSH along with cell viability could be further increased (Figures S1F and S1G) by treatment with an esterified analogue of GSH that is more permeable (23).

**Figure 1.**
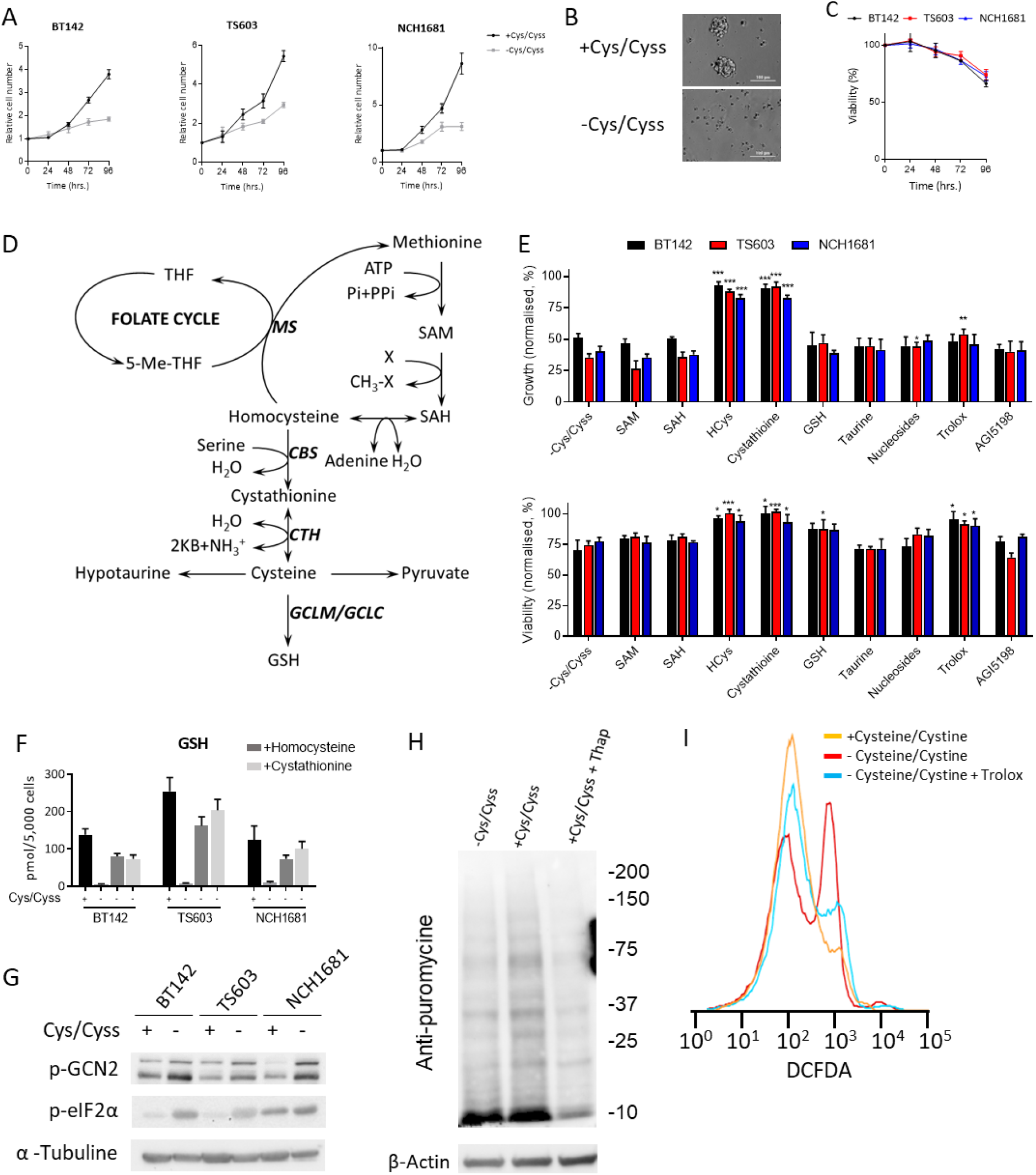
Cysteine and cystine deprivation halt growth and reduce viability in glioma through reduced protein translation and glutathione synthesis: (A) Growth rate of IDH1-mutant glioma cell lines under cysteine/cystine deprivation (Cys/Cyss) conditions over 96 hours. Data are shown as the ratio of viable cells to cells seeded (n = 4, data displayed as mean ± SD for each time point and condition). (B) Neurosphere formation after 96 hours of incubation in the same medium. BT142 cells displayed as a representative picture. (C) Viability after 96 hours in medium lacking cysteine and cystine as the percentage over the viability of cells in control conditions (n = 3, data displayed as mean ± SD for each time point and condition). (D) Metabolic pathways related to cysteine (THF, tetrahydrofolate; MS, methionine synthase; CBS, cystathionine beta-synthase; CTH, cystathionine gamma-lyase; GCLM, glutamate-cysteine ligase modulatory subunits; and GCLC, glutamate-cysteine ligase catalytic subunits). (E) Rescue experiments involving the addition of different metabolites and agents to cells grown without cysteine/cystine (data displayed as control-normalized growth [top panel] and viability [bottom panel] as the mean ± SD for n = 3; \**p* < 0.05, \*\**p* < 0.005, \*\*\**p* < 0.001, each condition was compared with the -Cys/Cyss values by using a two-tailed Student’s *t*-test). (F) Glutathione (GSH) levels in complete medium (+) and medium lacking cysteine/cystine (-) plus homocysteine and cystathionine (data displayed as the mean ± SD for n = 3). (G) Western blot of p-GCN2 and p-eIF2α in both complete and cysteine/cystine-free media for the 3 cell lines investigated. (H) Western blot shows puromycin-labeled proteins after 72 hours. Thapsigargin-treated samples are included as negative controls. BT142 cell line data is shown as a representative blot. (I) DCFDA assay as a marker of ROS production in glioma cell lines under cysteine/cystine deprivation and treated with 0.25 mM Trolox for 96 hours. BT142 cell line data is shown as a representative diagram.

Tumors have a higher proliferative rate, which translates into higher levels of oxidative stress (24), which can engage cancer cells in ROS-dependent death. We explored the mechanism of cell death by treating the cells with ferrostatin-1 (at cell seeding time), an inhibitor of ferroptosis (25), and then analyzing apoptotic markers (Figure S1I and S1J). Cell viability did not recover after treatment with ferrostatin 1 (Figure S1H), and an increase in apoptotic and preapoptotic cells was detected (Figure S1J) after 96 hours. Accordingly, the mechanism of cell death observed in these cell lines under cysteine/cystine depletion can be assigned to apoptosis, a process that can be triggered by high levels of ROS (26, 27). Thus, the antiproliferative effects observed in our study occurred in part from cells’ inability to synthesize cysteine for downstream protein synthesis, whereas the loss of viability was attributed to their inability to fight ROS.

### Global metabolic consequences of cysteine and cystine deprivation

Next, we explored the metabolic changes resulting from this nutrient deprivation condition after 48 hours (Figure 2A), before cellular viability is affected (Figure 1C); thus, we could analyze the metabolism without the potential confounding factors attributable to the mechanisms of cell death. The main metabolites affected were those directly related to the pathways that intrinsically involve cysteine/cystine, as well as those involved in related metabolic routes, such as the tricarboxylic acid (TCA) cycle (Figure 2A). In our metabolomic analysis, citrulline levels were also significantly lower after 48 hours of cysteine/cystine deprivation, but we could not detect any additional metabolites from the urea cycle of the arginine biosynthesis pathway. However, arginine levels in NCH1681 were significant higher under cysteine/cystine deprivation. To consider all these metabolic changes together, we performed a pathway analysis to identify the routes that were most affected (Figure 2B). We also evaluated the potential upregulation of enzymes connecting the methionine cycle with cysteine synthesis in order to compensate for the lack of cysteine/cystine. Expression of the first enzyme involved in the TS pathway, cystathionine beta-synthase (CBS) was not upregulated (Figure 2C), which reveals a metabolic disconnection between the methionine cycle and the TS pathway, even though the subsequent enzyme, cystathionine gamma-lyase (CTH), was upregulated.

**Figure 2.**
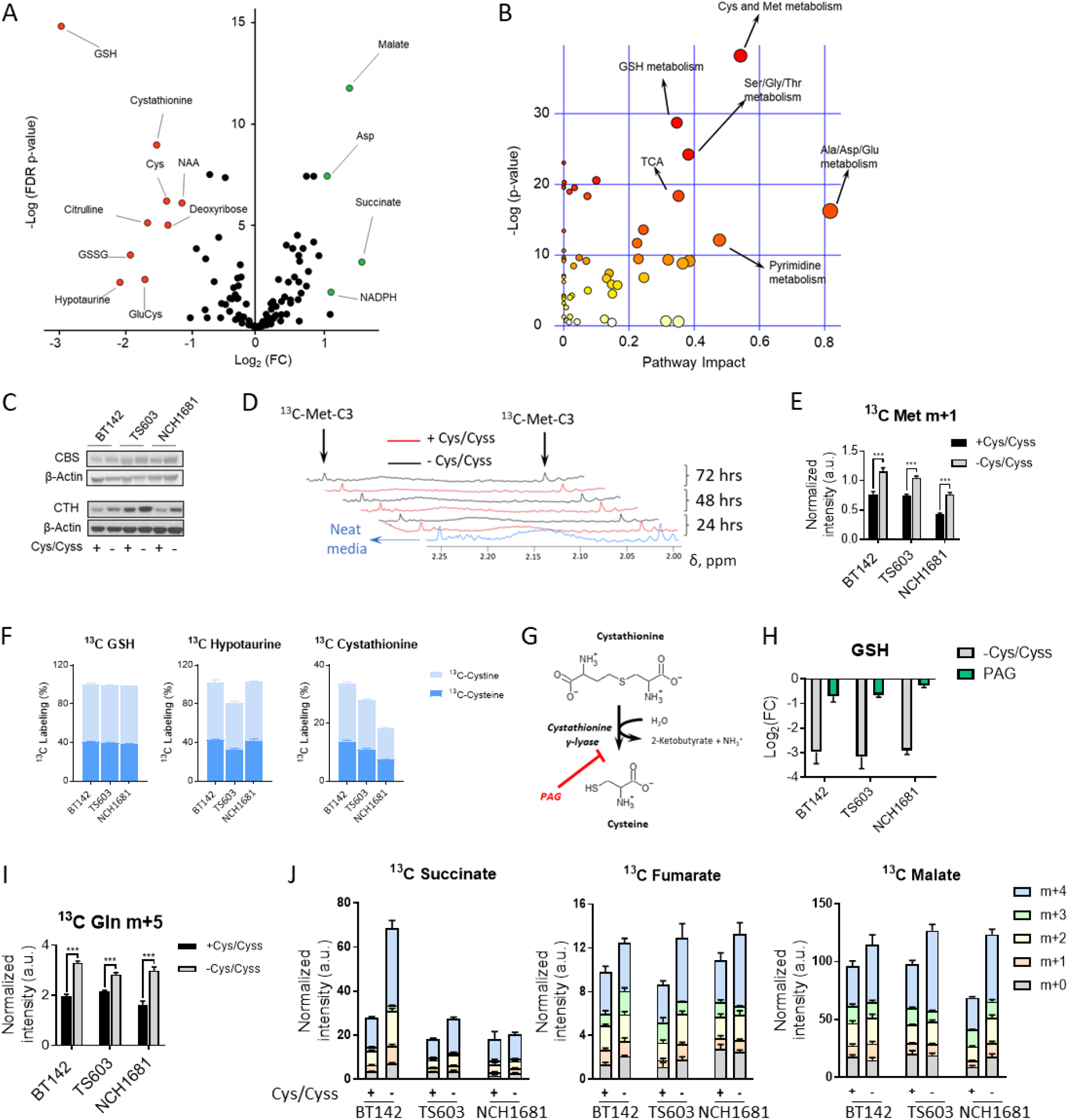
Global metabolic consequences of cysteine and cystine deprivation: (A) Volcano plot displaying the Log_2_(FC) vs Log(FDR *p* value) for all the metabolites identified by LC-MS. Metabolites highlighted in green (upregulated upon cysteine/cystine deprivation) or red (downregulated) have Log_2_(FC) > 1 or < −1 and an FDR *p* value < 0.05 (5 replicates per cell line and per condition). (B) Pathway analysis performed on the metabolites levels computed depicts the major pathways affected. (C) Western blot of cystathionine beta-synthase (CBS) and cystathionine gamma-lyase (CTH). (D) NMR spectra of media over time, including assignments for *methyl*-^13^C-methionine. Representative spectra shown as normalized intensities to the TSP signal (δ = 0.00 ppm) and cell number. (E) *Methyl*-^13^C-methionine levels in cells seeded in medium containing this tracer (data displayed as bar plots; mean ± SD, n = 3; \*\*\**p* < 0.001, two-tailed Student’s *t*-test). (F) ^13^C-tracing experiments displaying ^13^C-C3-cysteine and ^13^C-C3,3’-cystine incorporation into cystathionine, GSH, and hypotaurine (n=3 samples per cell line and ^13^C probe). These experiments were conducted separately for each tracer (^13^C-C3-cysteine and ^13^C-C3,3’-cystine). (G) Metabolic effect of propargylglycine (PAG) due to inhibition of the TS pathway, and (H) decrease of GSH levels (n = 5 samples per cell line) for both PAG treatment and cysteine/cystine deprivation displayed as Log_2_(FC). (I) ^13^C incorporation from ^13^C-U-glutamine into glutamine pools, and (J) TCA metabolites displayed as the contribution of each isotopologue to the total metabolite pool. ^13^C tracing experiments are displayed as bar plots; mean ± SD, n = 3 (\*\*\**p* < 0.001, two-tailed Student’s *t*-test).

We then explored whether methionine availability limited the cells’ ability to upregulate the TS pathway. We identified the ^13^C-*methyl* resonance signal from methionine in the media even after 72 hours, which indicated that methionine availability is not responsible for restricting cell growth even in the absence of cysteine/cystine (Figure 2D). Concurrently, we detected the accumulation of *methyl*-^13^C-labeled methionine in cysteine/cystine-deprived cells (Figure 2E). Methionine is a proteinogenic amino acid, and the downregulation of protein synthesis (Figures 1G, 1H, and S1E) likely increases the intracellular pool of methionine. Indeed, ^35^S-methionine/cysteine labelling followed by autoradiography (or other detection methods) can be used to assess the biosynthesis of proteins. The methyl group of methionine can also be transferred to nucleotides to contribute to the epigenetic landscape; lipids and proteins (28, 29) and the resulting metabolite, SAH, can be further transformed into homocysteine or be remethylated to generate methionine. Accordingly, we used metabolomics analysis to determine whether the total levels of metabolites involved in these related pathways would be affected by cysteine/cystine deprivation. Global profiling experiments using liquid chromatography–mass spectrometry (LC-MS) did not reveal any effect on the synthesis of methyl donors and folate metabolism (Figure S2B and S2C). Methyl donors were assessed by the ratio of SAM to SAH (Figure S2D) as an indicator of cellular methylation capacity (30). Additionally, we investigated the contribution of cysteine and cystine to derived metabolites through ^13^C-C3-cysteine and ^13^C-C3,3’-cystine labeling, respectively, and LC-MS experiments. After 48 hours, nearly 100% of GSH was derived from exogenous cysteine/cystine (Figure 2F). We also observed an active reverse reaction of CTH, which was inferred from the labelling of cystathionine from ^13^C-labeled cysteine and cystine, and the dependence of hypotaurine synthesis on exogenous cysteine/cystine. To determine the metabolic effects of inhibiting the TS pathway, we treated cells with 1 mM propargylglycine (PAG) (Figure 2G) for 48 hours. This treatment did not extensively affect the metabolic profiles of our cell lines (Figures S2E and S2F), although GSH levels were slightly lower but far from the 10-fold decrease (Figure 2H) obtained after cysteine/cystine deprivation. These analyses revealed that the TS pathway is not intensively active in these cells, which rely almost completely on exogenous cysteine/cystine for GSH synthesis. We incubated our cells in C3-^13^C-serine, which revealed a labeling of <1% in GSH in the presence of cysteine and cystine (Figure S2G) and a slight upregulation in media lacking both amino acids. Additionally, we observed an accumulation of m+5 glutamine (Figure 2I) along with an increased flux of glutamine toward the TCA cycle, as it is no longer needed for GSH synthesis (through glutamate conversion) or cystine import (31). Accordingly, glutamate-derived metabolites involved in the TCA cycle appear to be upregulated (Figure 2J). In contrast, nucleotide synthesis from glutamine was downregulated (Figure S2H), possibly due to halted cell proliferation, which decreased the demand for nucleotide biosynthesis.

### Redox homeostasis in glioma cell lines requires exogenous cystine and cysteine

IDH1-mutant enzyme utilizes NADPH to reduce α-ketoglutarate to D-2-hydroxyglutarate, which is also needed to regenerate GSH from GSH-disulfide (GSSG) catalyzed by glutathione reductase (Figure 3A). The ratio of NADP to NADPH, a marker of the antioxidation capacity of the cell (32), becomes imbalanced when cysteine availability is limited (Figure 3B). Additionally, we observed a significant reduction in the GSH:GSSG ratio (Figure 3C), although the availability of NADPH was higher in cysteine/cystine-starved cells. This decrease in the GSH:GSSG ratio indicated reduced glutathione reductase activity when glioma cells were grown in a cysteine/cystine-lacking medium, as the expression levels of the enzyme were maintained (Figure S3A).

**Figure 3.**
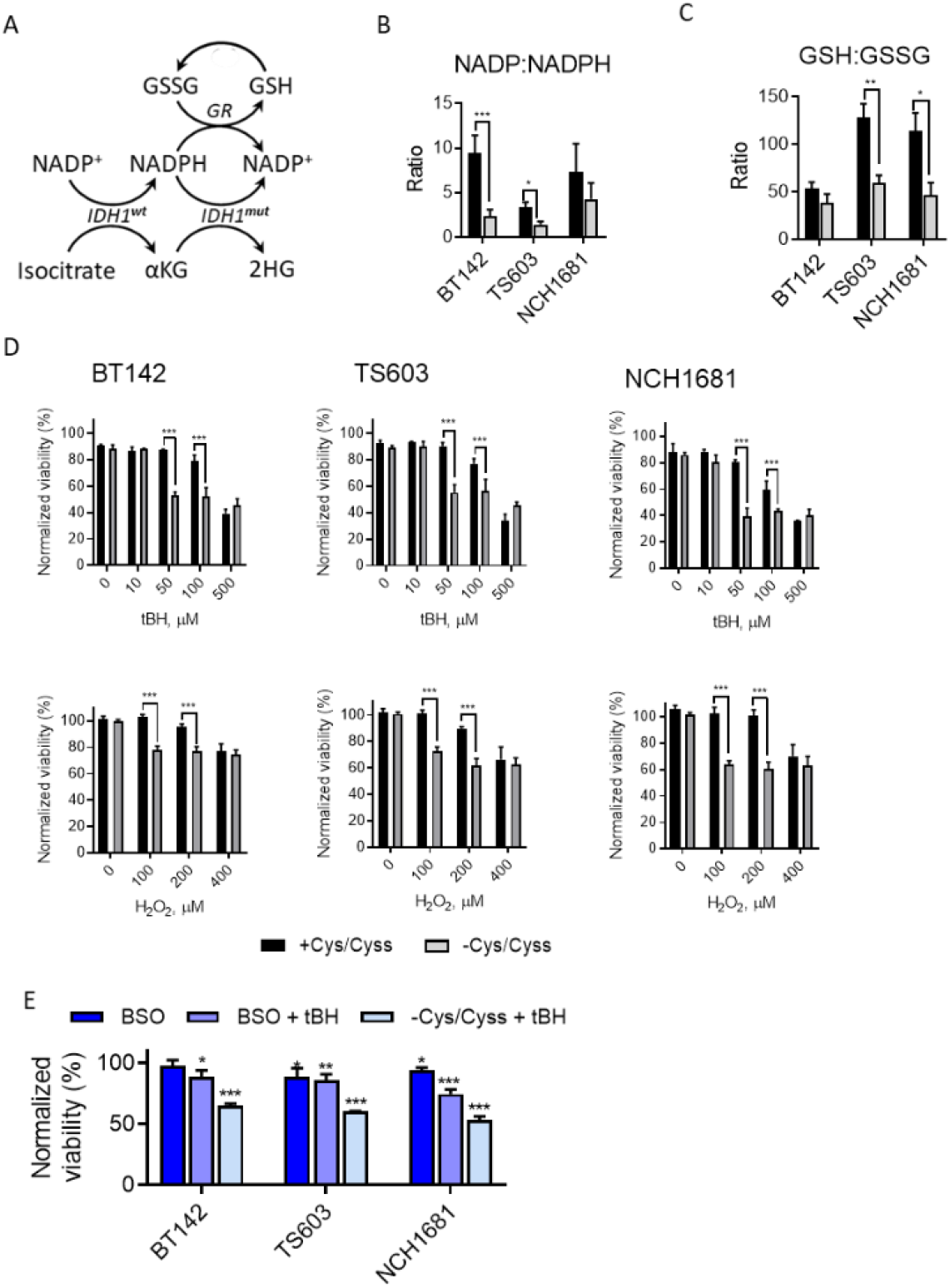
Redox homeostasis in glioma cell lines requires exogenous cystine and cysteine: (A) Simplified representation of the links between NADPH, IDH1 mutation, and GSH metabolism (GR, glutathione reductase). (B) Ratio of NADP and NADPH, and (C) GSH and GSSG grown in full medum (black) or cysteine/cystine-free medium (grey). Data are mean ± SD, n = 5 (\**p* < 0.05, \*\**p* < 0.005, \*\*\**p* < 0.001, two-tailed Student’s *t*-test). (D) Titration experiments involving tBH or H_2_O_2_ (data are mean ± SD, n = 3; \*\*\**p* < 0.001, 2-way ANOVA followed by Sidak’s multiple comparison test). (E) Effect of BSO plus tBH on viability. Data are mean ± SD of viability values normalized to the control conditions (full medium), n = 3; \**p* < 0.05, \*\**p* < 0.005, \*\*\**p* < 0.001, 2-way ANOVA followed by Dunnet’s multiple comparison test for control vs. all).

We also tested another main antioxidant system that was recently linked to tumor progression (33), thioredoxin, as a potential compensatory mechanism for GSH depletion; however, its expression levels remained unchanged when cells were deprived of cysteine (Figure S3B). These findings suggest that GSH depletion does not trigger the compensatory antioxidative mechanisms involving overexpression of thioredoxin in the time frame tested.

To exploit these findings therapeutically, we tested whether our nutrient restriction sensitizes these cancer cell lines to ROS and induces oxidative stress, as this approach has been shown to effectively induce the death of other types of cancer cells (34, 35). Thus, we conducted short titration experiments with tert-butyl hydroperoxide (tBH) and H_2_O_2_ (Figure 3D) by seeding the cells for 24 hours in the different media and then treating them for 6 hours with those ROS-inducing agents. This approach decreased cellular viability more intensively in cells grown for 24 hrs without cysteine/cystine. Indeed, 50 µM of tBH selectively induced cell death in 30%–50% of cells but did not affect the viability of the cells grown in full medium. Likewise, 100 µM of H_2_O_2_ had the same deleterious effect. That ROS-induced cell death could be partially reversed by adding Trolox or GSH (Figure S3C). Traditionally, buthionine sulphoximine (BSO) has been the drug of choice to target GSH synthesis and increase the sensitivity of cancer cells to oxidative stress; however, that approach has not been effective in some cases (31), including gliomas as a single agent (36). We observed similar results for the BT142 cell line, in which BSO as a single agent did not reduce cellular viability (Figure 3E), and the effect was much lower than that achieved by our approach. Similarly, nutrient limitation had a more intense effect on TS603 and NCH1681 cells because, unlike BSO, nutrient deprivation also affects the cells’ ability to regenerate GSH through glutathione reductase activity (37). We also tested whether cysteine/cystine deprivation synergizes with therapeutic agents, such as temozolomide (TMZ), that are typically part of the standard care for glioma therapy. The combination of nutrient restriction and TMZ (Figure S3E) improved the effect of TMZ for doses of 0.01 and 0.1 mM, but that decrease in viability was not more intense than the one generated by cysteine/cystine deprivation alone. At 1 mM TMZ, the cells experienced a more intense effect, but it was not enhanced by cysteine/cystine deprivation.

### Cysteine/cystine-free diet reduces the levels of glutathione temporally in a mouse model of glioma and increases survival

To test the effect of cysteine/cystine-deprivation on survival, we conducted an *in vivo* experiment in which we injected mice intracranially with NCH1681 cells. Twelve days later, we randomized the mice into 2 groups and provided a cysteine/cystine-free or a control diet the following day. We then monitored the plasma levels of the main metabolites related to cysteine, as potential biomarkers of treatment response at 16, 35, 44, and 55 days after intracranial injection. For each time point, metabolite levels were normalized to those computed in the control group. GSH and cystine levels in plasma were significantly lower in the cysteine/cystine-free diet group 23 days after initiating the nutritional intervention (Figure 4A). We also performed a ^13^C-tracing experiment *in vivo* at the study end point by injecting ^13^C-U-glutamine into the tail vein and tracking the incorporation of this metabolite into glutamate, glutamine, and GSH in order to evaluate the ability of tumor cells to synthetize GSH *de novo*. We observed active *de novo* synthesis of GSH in the cysteine/cystine-free diet group, similar to that detected in the control group (Figure 4B). Moreover, levels of ^13^C-labeled m+5 glutamine were higher in the tumor region for both cohorts, as previously reported (38), and lower glutamate m+5 levels correlated more strongly with the cysteine/cystine-free diet. GSH m+5 levels were higher than those detected in the respective contralateral regions, in accordance with a higher need for antioxidant capacity in tumor cells (33, 39). Levels of ^13^C-labeled GSH in tumor tissue under both diets were consistent with the recovery circulating levels of cystine in plasma (Figure 4A), indicating a systemic response to the nutritional intervention. Additional metabolites related to the cysteine pathway but diverted from the synthesis of GSH, such as taurine and hypotaurine, showed a different trend than those of cystine and GSH; i.e., their levels diminished over time under the cysteine/cystine-free diet (Figure S4). This temporary reduction in the main antioxidant source translated to increased median survival of the mice by 26 days and enhanced oxidative stress in the tumor tissue of mice given the cysteine/cystine-free diet (Figures 4C and D).

**Figure 4.**
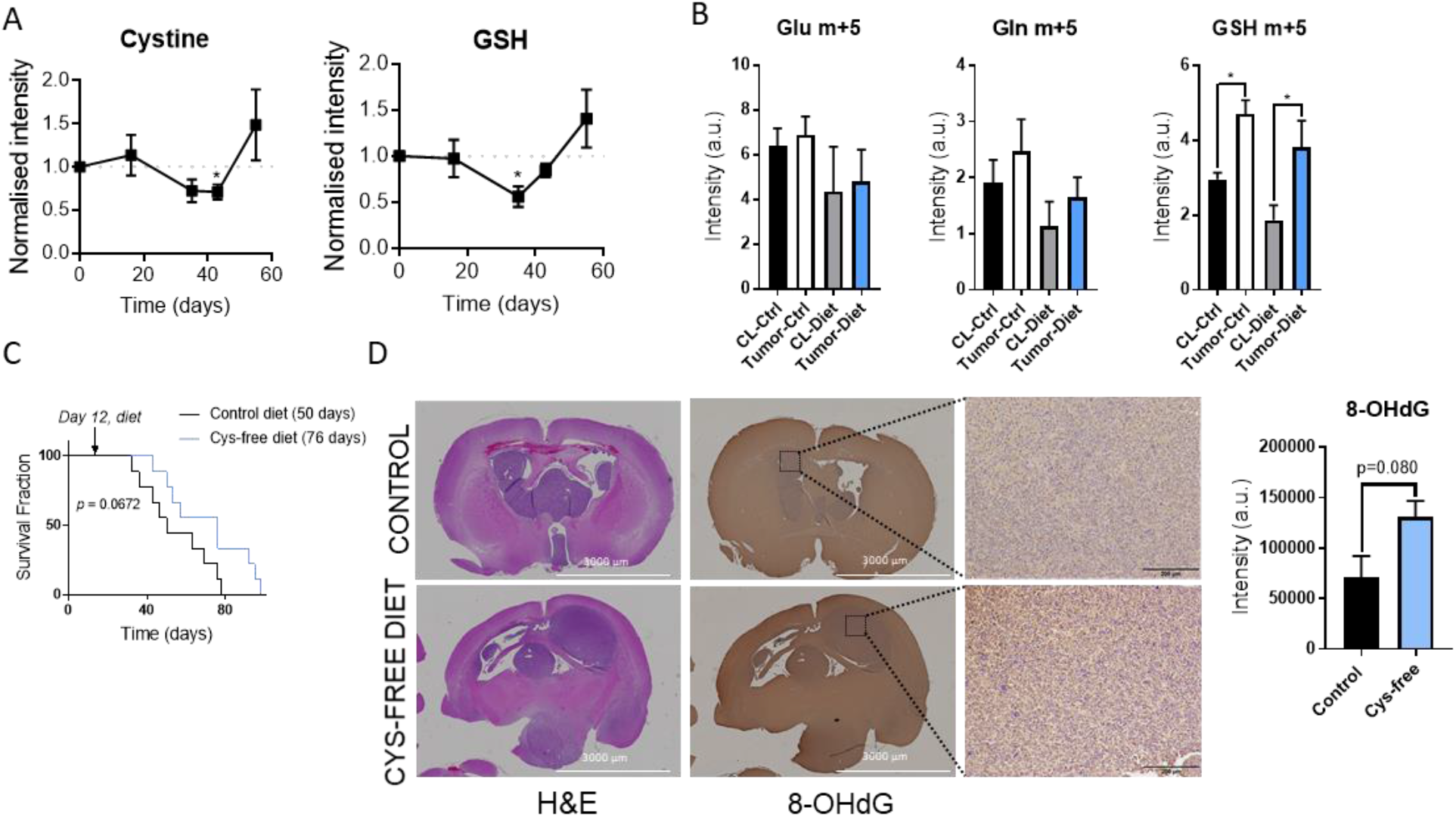
Cysteine/cystine-free diet reduces the level of glutathione temporally in a mouse model of glioma and increases survival: (A) Plasma levels of GSH and cystine normalized to those computed for the control group (dotted line, reference values of the control group; metabolite levels for the cysteine/cystine-free diet group are shown as mean ± SEM; n = 5; \**p* < 0.05 by two-tailed Student’s *t*-test with Welch’s correction). (B) Glutamate, glutamine, and GSH m+5 levels in tumor tissue and contralateral (CL) regions of mice fed with the control (Ctrl) and cysteine/cystine-free diets resulting from bolus injection of ^13^C-U-glutamine. Data are mean ± SEM of the normalized intensities to tissue weight, n = 3–5. \**p* < 0.05 by a one-way ANOVA followed by Tukey’s multiple comparisons test. (C) Survival analysis of intracranial glioma mouse model under both diets (n = 9, *p* value obtained from a Mantel-Cox test, and median survival displayed between brackets). (D) Immunohistochemical analysis of hematoxylin and eosin (H&E)- and 8-hydroxydeoxyguanosine (8-OHdG)-stained tissues, including quantification of 8-OHdG (bars display mean ± SD, n = 3 mice).

## Discussion

We have shown how dietary limitation of the nonessential amino acid cysteine has an antiproliferative effect and can reshape the metabolic landscape of cancer cells. Although the deprivation of cysteine was recently shown to have a deleterious effect on tumor growth (40), the metabolic consequences triggered by this situation were unexplored. Here, we provided a broad description of the perturbations occurring in the metabolic pathways of glioma cells, highlighting the absence of an active TS pathway and offering a supplementary window of intervention during which to reduce the tumor’s ability to fight oxidative stress. It has been reported that limiting the exogenous contribution of cysteine is a potential treatment for prostate, pancreas, and breast cancers (41, 42); renal cell carcinomas (43); and leukemia (44).

The system known to divert homocysteine into cysteine involves the upregulation of CBS activity together with increased levels of SAM (45), which drives homocysteine into cystathionine, an irreversible reaction. Homocysteine levels must be maintained at low concentrations because an excess of this metabolite can increase ROS (46); accordingly, cells have developed clearing systems by remethylating it, yielding methionine, or by launching the TS pathway. Recently, a disconnection between both pathways was reported in cancer cell lines because the cells are unable to modulate the methylating enzyme (40), glycine N-methyltransferase, which was resolved by ectopic overexpression of this enzyme. The methylation capacity of the glioma cell lines utilized in our study did not change a result of cysteine-limited conditions (Figure S2C). Rather, the constraint lies in the ability of the cells to divert the methionine cycle and generate downstream metabolites rather than upregulate enzymes involved in this route, because addition of the substrates for CBS and CTH recovered the cells’ growth and viability (Figure 1E).

We detected incorporation of the ^13^C label from cysteine/cystine to cystathionine as well as overexpression of CTH after cysteine/cystine deprivation; thus, this result might suggest that cystathionine can act as a reservoir of cysteine. Then, in nutrient-limited conditions, cystathionine can be broken down to yield cysteine. The disconnection between the TS pathway and the methionine cycle was reported recently in similar experiments that found that after addition of either cystathionine or homocysteine, both proliferation and recovery were restored, but not when cells were supplemented with methionine in a cysteine-deficient medium (47).

It is not known why some cancers can upregulate the TS pathway and do not rely on the uptake of cystine and cysteine (16). Cysteine/cystine deprivation caused a 10-fold reduction in GSH levels in our cell lines, thereby hypersensitizing glioma cells to ROS and revealing a high dependence on the exogenous supply of this amino acid. A recent investigation also showed how tumors that have large antioxidant capacity require an exogenous supply of nonessential amino acids for proliferation (48). Targeting GSH synthesis has been a recurrent therapeutic strategy (41), mainly to sensitize tumor cells in combinatorial therapies (49-51). That approach has also been tested in IDH1-mutant cancers (36), although in the present work we observed how IDH1-wildtype gliomas were as sensitive as mutant gliomas to cysteine/cystine deprivation and that inhibiting IDH1-mutant activity did not affect the response of those cells to this approach.

Tumor cells display enhanced oxidative stress, which is compensated for by a larger antioxidant capacity (11, 39). Therefore, reducing the ability of tumor cells to fight against ROS can be an efficient strategy for treating cancer. From a pharmacologic point of view, brain tumors are challenging because of the selective permeability of the blood-brain barrier. Through our strategy, we could target the antioxidant capacity of tumor cells without using any drug therapy. Herein, we show that a cysteine-deprivation diet creates a window of therapeutic opportunity by decreasing GSH and its precursor (cystine/cysteine) at the systemic level (Figures 4A and S4), which could then be utilized in combination with a ROS-inducing agent. Although the plasma levels of these metabolites (GSH and cystine) were recovered afterwards, those of hypotaurine and taurine (Figure S4), which are generated through the cysteine dioxygenase pathway, were not recovered. Nevertheless, homocysteine and methionine levels did not change significantly due to the diet (Figure S4). The selective recovery of metabolites related to the antioxidant response might indicate an enhanced demand for ROS-buffering agents due to the increased oxidative stress generated by the diet. Nutritional interventions have been reported recently to be efficient strategies for tumor treatment in animal models (6), including a cysteine-limitation strategy in a model of pancreatic cancer (52). Our approach can be used clinically when it is necessary to limit the antioxidant capacity to enhance the effect of a secondary drug in a combinatorial treatment or when the resistance mechanism to a treatment depends on the ability of the cells to upregulate GSH metabolism.

### Limitations of the study

Our investigation focused on IDH1-mutant gliomas because they have been traditionally described as being more vulnerable to oxidative stress, although this specificity was contrasted through treatment with AGI5198 and by including GSC923 and GSC827 cell lines (IDH1-wildtype) in the study. Nevertheless, future investigations involving a mouse model of IDH1-wildtype glioma will provide additional evidence for our findings. The *in vivo* experiment correlated the results from time series analysis of plasma with those obtained from tissue at end point; therefore, a longitudinal analysis of tumor tissue after nutritional intervention would add more value to our interpretation. This approach would require GSH to be monitored in tissue through magnetic resonance spectroscopy or by euthanizing the animals at different time points to extract the metabolites for analysis.

## Materials and Methods

### Cell culture

Cell lines utilized in this investigation: TS603 (53) was a gift of T. A. Chan (Memorial Sloan-Kettering Cancer Center, USA); NCH1681 (54) was provided by C. Herold-Mende (University Hospital Heidelberg, Germany); BT142 was purchased from ATCC^®^ (ACS-1018™); and GSC923 and GSC827 were generated at the Neuro-Oncology Branch of the National Institutes of Health (Bethesda, USA). All cell lines were authenticated using DNA sequencing and the Illumina platform to detect the glioma methylome and the 1p/19q co-deletion.

All cell lines were grown in DMEM:F12 medium supplemented with antibiotics (penicillin-streptomycin), 1% N2 growth supplement, heparin sulfate (2 µg/mL), EGF (20 ng/mL), and FGF (20 ng/mL). For experiments involving seeding the cells in medium containing ^13^C tracers or lacking cysteine/cystine, neurospheres grown in regular medium were collected, centrifuged, and resuspended in phosphate-buffered saline (PBS). Cells were then spun down, the supernatant was discarded, the pellet was resuspended again in PBS, and neurospheres were mechanically disaggregated for cell counting and subsequently seeded in the corresponding medium. Medium lacking cysteine and cystine was provided by Thermo Fisher upon request. C3-^13^C-cysteine (Cambridge Isotopes Laboratories, CLM-1868) and C3,3’-^13^C-cystine (Cambridge Isotopes Laboratories, CLM-520) were utilized for ^13^C tracing. Experiments involving *methyl*-^13^C-methionine (Cambridge Isotopes Laboratories, CLM-206-1), C3-^13^C-serine (Cambridge Isotopes Laboratories, CLM-1572), and U-^13^C-glutamine (Cambridge Isotopes Laboratories, CLM-1822) were performed in DMEM:F12 using in-house made medium that lacked the metabolites utilized as ^13^C tracers. We created 20x–100,000x solutions (according to solubility) of all the components described in the DMEM:F12 formulation. For compounds that required a non-neutral pH to be soluble in water, a 1N solution of NaOH or HCl was used to titrate the solutions. Components of the medium were added to reach the final concentrations described in the commercial version of DMEM:F12, but linoleic acid was included directly from the vial provided by the vendor. Distilled water was added to obtain the final volume desired. This medium was subsequently filtered, and the supplements described above were added, and it was filtered again. All the ^13^C probes were utilized in the same concentrations as those specified in the DMEM:F12 commercial formulation.

### Treatments and proliferation assays

A total of 50,000 to 100,000 cells/well were seeded in untreated 48-well plates with 0.75 mL of culture medium in triplicate (all experiments were repeated at least twice). After treatment, neurospheres were mechanically disaggregated and counted using the Vi-CELL XR cell counter (Beckman Coulter, USA). Glutathione was added at a final concentration of 0.5 mM. S-adenosylmethionine, homocysteine, cystathionine, and taurine were dissolved in PBS, filtered, and added to a final concentration of 0.1 mM. Nucleosides (100x; Millipore Sigma ES-008-D) were added directly to the cultured cells after filtration to a 1x final concentration. AGI5198 was dissolved in DMSO and used in a 10 µM final concentration. BSO and Trolox were dissolved in PBS and DMSO, respectively, at 0.25 mM final concentrations. Ferrostatin 1 was utilized at a final concentration of 2 µM. H_2_O_2_ and tBH were dissolved in PBS, filtered, and added to the wells to reach the final concentrations specified in the graphs. Puromycin assay for detecting protein synthesis was performed as described in (18).

### DCFDA assay

Experiments were performed employing the DCFDA/H_2_DCFDA-Cellular ROS Assay Kit (Abcam, ab113851) according to the vendor’s protocol and analyzed with a Sony SA3800 580 spectral analyzer. Data were processed and plots were generated with FlowJo 10.6 (BD Biosciences).

### Apoptosis assay

Experiments were performed using the 7AAD Annexin V Apoptosis Detection Kit (Abcam, ab214663) according to the vendor’s protocol and analyzed with a FACSCalibur (Becton Dickinson, Franklin Lakes, NJ, USA). Data were processed and plots were generated with FlowJo 10.6 (BD Biosciences).

### Sample preparations for metabolomics

Cells were collected by centrifugation and washed twice with PBS, and the resulting pellet was stored at −80°C until extraction. For extraction, cell pellets were thawed on ice and lysed by 3 cycles of freeze-thawing, including a 5-min sonication process in an ice-water bath during the thawing step. Then 40 μL of the homogenate were put aside for protein quantification by the Bradford method for further normalization of the metabolite levels. Next, metabolites were extracted by mixing the lysate with methanol:chloroform at a final ratio of 1:2:2 (water:methanol:chloroform), thoroughly vortexed, and incubated in ice with agitation for 10 minutes. Then samples were centrifuged at 12,000 rpm for 20 min at 4°C. The two resulting phases (upper aqueous polar and lower organic lipid) were separated, and the protein interface was discarded. Polar extracts were dried under a stream of N_2_ and stored at −80°C until metabolomics analysis was performed. Tumor tissue was first weighed while frozen for normalization purposes and subsequently homogenized using a bullet blender homogenizer in the same solvent mixture described above and further processed in the same way as cell extracts. Cell culture medium was collected after centrifugation of cells (300 rcf, 5 min), extracted in ice-cold methanol, dried under N_2_, and resuspended in 180 μL of phosphate buffer (pH 7; 100 mM) in D_2_O (containing d-TSP) and 1% NaN_3_ for nuclear magnetic resonance (NMR) spectroscopic analysis.

### NMR spectral acquisition and processing

All spectra were acquired at 25°C on a Bruker Avance III 600 MHz spectrometer (Structural Biophysics Laboratory, NCI, Frederick, MD, USA) equipped with a cryogenically cooled probe. Single-pulse ^1^H NMR experiments were performed using the noesygppr1d (TopSpin 3.5, Bruker Biospin) pulse sequence for water suppression. For each spectrum, 128 scans were acquired, with a relaxation delay of 3 s, a spectral width of 10,800 Hz, and a time domain of 32K points. Spectra were referenced to the TSP internal standard signal (s, δ = 0.00 ppm), zero-filled to 64K points, phased, and baseline-corrected using ACD Labs Spectrus Processor 2016, and an exponential line-broadening function of 0.30 Hz was applied. For quantification, ^1^H NMR resonance signals were normalized to the TSP singlet located at 0.00 ppm and corrected to either the total protein content as obtained from the Bradford assay, cell number, or tissue weight.

### LC-MS acquisition and processing

LC-MS analysis was conducted with an Agilent 6545 MS combined with a 1290 Infinity II UHPLC system. Only LC-MS-grade solvents and additives purchased from Covachem, LLC (Loves Park, IL, USA) were used to prepare mobile phases and wash solutions. Wash cycles consisting of a strong wash (50% methanol, 25% isopropanol, and 25% water), weak wash (90% acetonitrile and 10% water), and seal wash (10% isopropanol and 90% water) were implemented to eliminate carryover between injections. Analytes were injected (8 µL) and resolved using an Infinity 1290 in-line filter combined with an AdvanceBio Glycan Map 2.1 × 100 mm, 2.7 µm column (Agilent Technologies, Wilmington, DE, USA) set at 35°C. The solvent buffers, consisting of mobile phase A (10 mM ammonium acetate in 88% water and 12% acetonitrile) and mobile phase B (10 mM ammonium acetate in 90% acetonitrile), were initially titrated with formic acid and ammonium hydroxide to pH 6.85. The linear gradient was executed at a flow rate of 0.3 mL/min, as follows: 100% B, 0.5 min; 95% B, 2.0 min; 60% B, 3.0 min; 35% B, 5 min; hold 0.25 min; 0% B, 6 min; hold 0.5 min; 100% B, 7.8 min. The mass analyzer acquisition conditions were as follows: drying gas temperature 250°C, sheath gas temperature 325°C, nebulizer 45 psig, skimmer 50 V, and octopole radio frequency 750 V. Mass spectra were acquired at 3.0 spectra/s in negative electrospray ionization (ESI) mode for a mass range from 72 to 1200 m/z using a voltage gradient of capillary 3000 V, nozzle 2000 V, and fragmentor 80 V.

Prior to preprocessing the datasets, pooled quality control samples were inspected for consistency of retention-time (RT) shifts and signal degradation. Following acquisition, m/z spectra binning was performed by partitioning the m/z vs. RT matrices into fixed width using an Agilent Masshunter Profinder B.08.00. Bins were manually inspected to confirm consistent, reproducible integration for all analytes of interest across all samples. Target extraction of precursor m/z was performed using an in-house compound library. Ion selection and alignment parameters were restricted to proton loss (H-) only, 5.0 mDa mass range, and RT ± 0.4 min. Following preprocessing, the ion areas were reported for each sample and corrected to the area of sample-specific internal standard, p-nitrobenzoate (added at 90 pmol/per sample during preparation).

The same acquisition procedure was followed for the ^13^C isotopically labeled samples. After alignment and identification of analytes of interest retention times a PCDL card was constructed using PCDL Manager B.07.00 (Agilent). The chromatograms were introduced into the Agilent MassHunter Profinder B.08.00, and the PCDL card was used in the Batch Isotopologue Extraction routine with the following parameters: 99% ^13^C labeling, 20% peak height ion abundance criteria, mass tolerance of ±15 ppm + 2 mDa with a threshold of 250 counts for the anchor and 1000 counts for the sum of the ion heights with a minimum correlation coefficient greater than 0.5. The corrected and raw intensity and percentages of isotopologues of the analytes of interest were obtained. Metabolite levels obtained from the LC-MS analysis described above are displayed either as the percentage of the specific isotopologues over the total pool of the metabolite for ^13^C tracing experiments or as protein/tissue-normalized intensities.

### Metabolomic analyses

Volcano plots were generated from the datasets obtained from the LC-MS analysis of the polar extracts of the 3 cell lines. Samples were labeled as control or cysteine/cystine-free. Then *p* values obtained by a *t*-test followed by Welch correction were adjusted for multiple comparisons by the false discovery rate (FDR) method. In addition, fold changes (FC) were computed for each metabolite to generate the plot, including -Log(FDR-corrected *p* values) vs Log_2_(FC). Thresholds used to highlight dysregulated variables were Log_2_(FC) > 1 or < −1 and FDR-corrected *p* values < 0.05. Volcano plots and PCA data were obtained by MetaboAnalyst 4.0 (55).

### Western blots

Samples were harvested and centrifuged at 300 x *g* for 3 min, washed with PBS, and lysed in a solution of the radioimmunoprecipitation (RIPA) buffer system (Santa Cruz Biotechnology, USA). After a 30-min incubation period and a brief sonication on ice, lysates were centrifuged at 14,000 x *g* for 10 min at 4°C. Total protein concentration of supernatants was then determined by BCA assay (Thermo Scientific). Samples were prepared for electrophoresis as 20-50 µg of protein, Laemmli sample buffer, and dithiothreitol (DTT). Protein samples were boiled at 95°C for 5 min and loaded into either 8-16% or 10% Mini-Protean TGX Precast Gels for separation by electrophoresis. Then proteins were transferred to nitrocellulose membranes using the Trans-Blot Turbo system. Blots were blocked for 1 hour at room temperature with 5% fat-free milk or milk-free blocking buffer (for analysis of phosphorylated proteins) and incubated overnight at 4°C with primary antibodies for p-GCN2 (Abcam, ab75836), p-eIF2α (Cell Signaling, #9721), α-tubuline (Cell Signaling, #2144), β-actin (Abcam, ab8227), cystathionine β-synthase (Abcam, ab135626), cystathioninase (Cell Signaling, #30068), glutamate-cysteine ligase regulatory subunit (Proteintech, 14241-1-AP), puromycin (EMD Millipore, MABE343), MTHFD1 (Proteintech, 10794-1-AP), and glutathione reductase (Abcam, ab16801). Subsequently, the membrane was washed with Tris-buffered saline + Tween-20, incubated with horseradish peroxidase (HRP)-conjugated secondary antibodies for 1 h at room temperature and treated with Clarity Max Western ECL substrate. Blots were imaged on the Chemidoc MP imaging system (BioRad).

### GSH quantification

GSH levels were computed using the GSH-Glo™ Glutathione Assay (Promega, USA) according to the protocol provided by the vendor. Then 24-well plates were seeded at 200,000–300,000 cells in 2 mL for 4 days. To normalize GSH levels, 0.5 mL was taken from each well prior to performing the assay, and the cells were counted by a ViCell XR automatic cell counter.

### Animal studies

The intracranial orthotopic mouse model harboring the IDH1-mutant glioma cell line NCH1681 was established according to approved animal study proposal NOB-008 by the National Cancer Institute−Animal Use and Care Committee. Briefly, cells were harvested, washed with PBS, and counted. The resulting pellet was resuspended in Hank’s Balanced Salt Solution, and 5 μL of the cell suspension were injected stereotactically into the striatum of 6–8-week-old female severe combined immunodeficient (SCID) mice (Charles River Frederick Research Model Facility) using a stereotactic device. Before the diet was initiated, mice were randomized and split into the control and the diet groups. The number of animals per group was determined by using G*Power 2. The sample size (number of animals) was computed via a priori methods of calculation, assuming an alpha error probability of 0.05, power level of 0.95, and difference in mean survival between groups of 9 days. Neurological symptoms of mice were monitored daily to assess tumor growth; specifically, an external independent researcher assessed the health of the mice twice a day without previous knowledge of the experiment (blinded to the treatment). Once this researcher determined that a mouse was reaching end point, in view of the symptomatology, that mouse was euthanized. Symptoms include animal experiencing rapid weight loss (>15%, monitored daily), debilitating diarrhea, rough hair coat, hunched posture, labored breathing, lethargy, persistent recumbence, significantly abnormal neurological signs, bleeding from any orifice, self-induced trauma, impaired mobility, moribund, or otherwise unable to obtain food or water. To compare survival curves, the log-rank (Mantel–Cox) test was used (GraphPad Prism 7.05). Diets supplied to the mice were A05080217 (L-amino acid rodent diet without added cystine) and A18110101 (L-amino acid diet with 2 g L-cysteine and 2 g L-cystine per kg) from Research Diets Inc. (New Brunswick, NJ, USA). Prior to initiation of the experiment, control mice were fed with both diets to check whether there was active ingestion of food. Accordingly, food and animal weight were monitored twice a week, and stool was examined visually for incorporation of food dyes.

### ^13^C-tracing *in vivo*

^13^C tracing experiments *in vivo* were performed as previously described (56-58) on the same mice utilized for survival analysis. ^13^C-U-glutamine was prepared as a 36.2 mg/mL stock solution in sterile PBS and injected (200 μL, 7.24 mg) into the tail vein at 15-min intervals for 3 times (total = 142 μmol) just prior to mice reaching end point. Mice were euthanized 15 min after the last injection (45 min from the first injection). Tumors were separated from the brain, then both tumor and normal tissues were gently blotted by rapidly tapping the tissue onto a cloth and were immediately flash-frozen in liquid nitrogen.

### Plasma analysis

Blood was collected from the tail vein of the mice in lithium-heparin collection tubes (Sarstedt, #41.1393.105). Approximately 35 µL of blood was centrifuged, according to the tube manufacturer’s instructions, at 4°C, and the clear plasma fraction was transferred to a clean microtube. Subsequently, plasma was extracted in a water:methanol:chloroform mixture, centrifuged for 20 min at 4°C and 13,000 rpm, and the resulting upper hydrophilic phase was then transferred to a clean vial and dried under a stream of N_2_. Dried sediments were resuspended in methanol and injected into the LC-MS system for global profiling.

### Immunohistochemistry

When mice reached end point, tissue was collected and stored in paraformaldehyde at 4°C. Tissue was submitted to Histoserv, Inc. for analysis together with the antibodies. Slides were deparaffinized and hydrated through graded alcohols to distilled water, followed by antigen retrieval. They were then blocked with hydrogen peroxide and a blocking serum. Next, the slides were incubated with the primary antibody, a secondary antibody, and HRP-conjugated streptavidin. Finally, the slides were developed using 3,3’-diaminobenzidine and counterstained with hematoxylin. All of the incubations were carried out at room temperature, with Tris-buffered saline + Tween-20 used as a washing buffer. Immunostaining was quantified using ImageJ software.

### Statistical analysis and graphs

Statistical significance was computed with R or Prism GraphPad 7.05. Multivariate analysis was performed using MetaboAnalyst 4.0 (59) and in-house R scripts. For each figure, “n” refers to the number of biological replicates. Exact p-values for those experiments in which statistical significance was assessed can be found in the supplementary files.

## Acknowledgments

We thank Hua Song and Wei Zhang for their help with the animal work.

## Supplementary Materials

**Figure S1:**
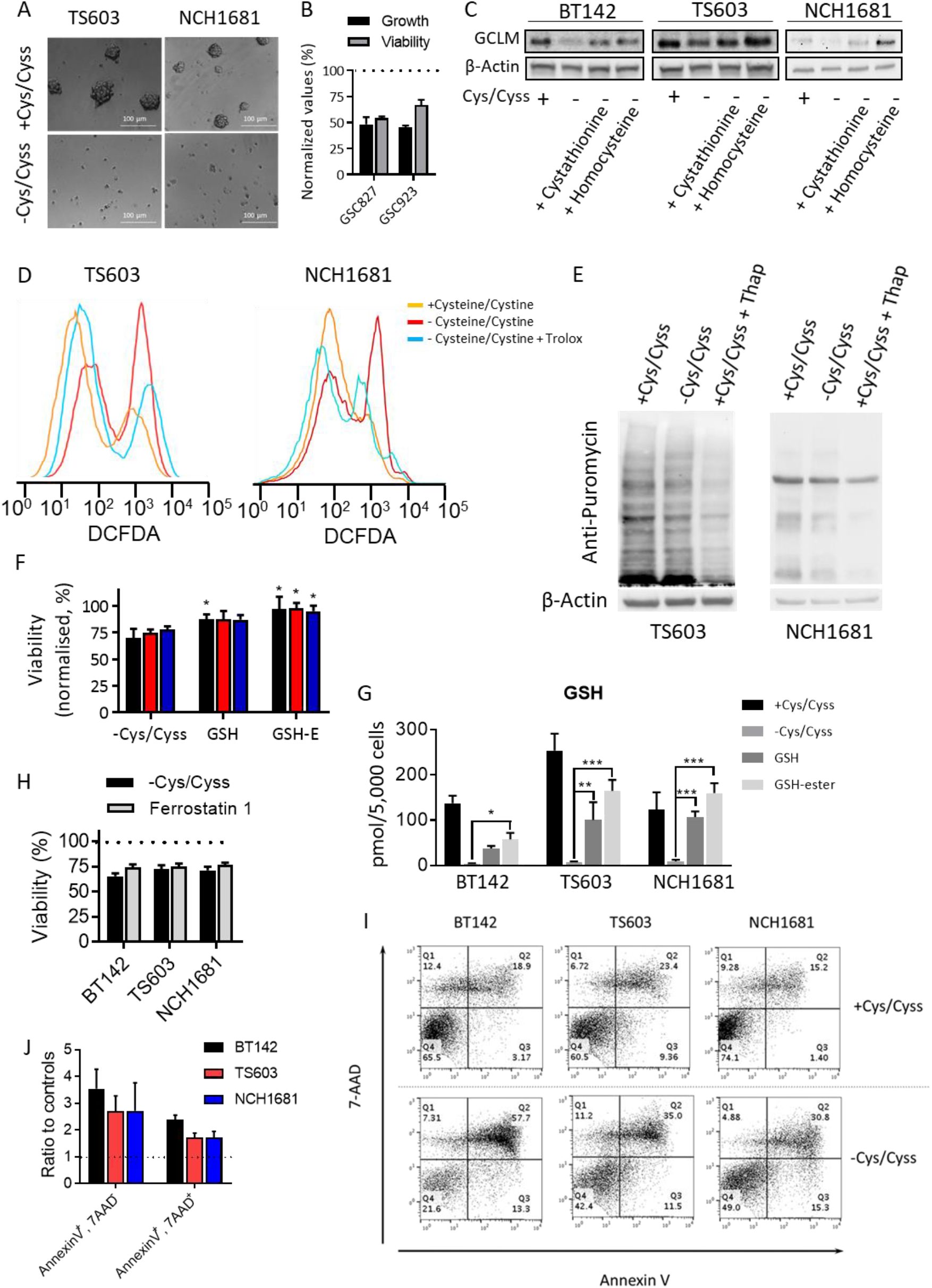
(A) Sphere formation of glioma cell lines after 96 hours of growth in medium with/out cysteine/cystine. Representative pictures are shown. (B) Growth and viability of IDH1-wildtype glioma cell lines under cysteine/cystine-deprivation for 96 hours (n = 3, bar plots show mean ± SD normalized to results from experiments performed in full medium). (C) Western blots of glutamate cysteine ligase modulatory subunit (GCLM) for 3 IDH1-mutant glioma cell lines grown in medium without cysteine/cystine and treated with 0.1 mM cystathionine or homocysteine. (D) DCFDA area as a marker of ROS in glioma cell lines under cysteine/cystine deprivation and treated with 0.25 mM Trolox. (E) Puromycin-labeled proteins detected by western blotting. (F) Viability of glioma cell lines in cysteine/cystine-lacking medium supplemented with either GSH or GSH-ethyl ester (mean ± SD for n = 3; \**p* < 0.05 versus control by two-tailed Student’s *t*-test) and (G) quantification of intracellular GSH (mean ± SD for n = 3; *, *p* < 0.05; \*\**p* < 0.005; \*\*\**p* < 0.001 from a one-way ANOVA followed by Tukey’s multiple comparison test for cells grown in cysteine/cystine-free medium vs those grown in the same medium plus either GSH or GSH ethyl ester). (H) Viability of glioma cell lines after 96 hours in medium lacking cysteine/cystine and in the same medium but treated with 2 µM ferrostatin 1 (n = 3, bar plots show mean ± SD normalized to results from experiments performed in full medium). (I) Apoptosis detection assay diagrams at 96 hours and (J) the corresponding quantification of the number of cells assigned to early apoptosis and apoptosis (n = 3, bar plots show mean ± SD normalized to results from experiments performed in full medium).

**Figure S2:**
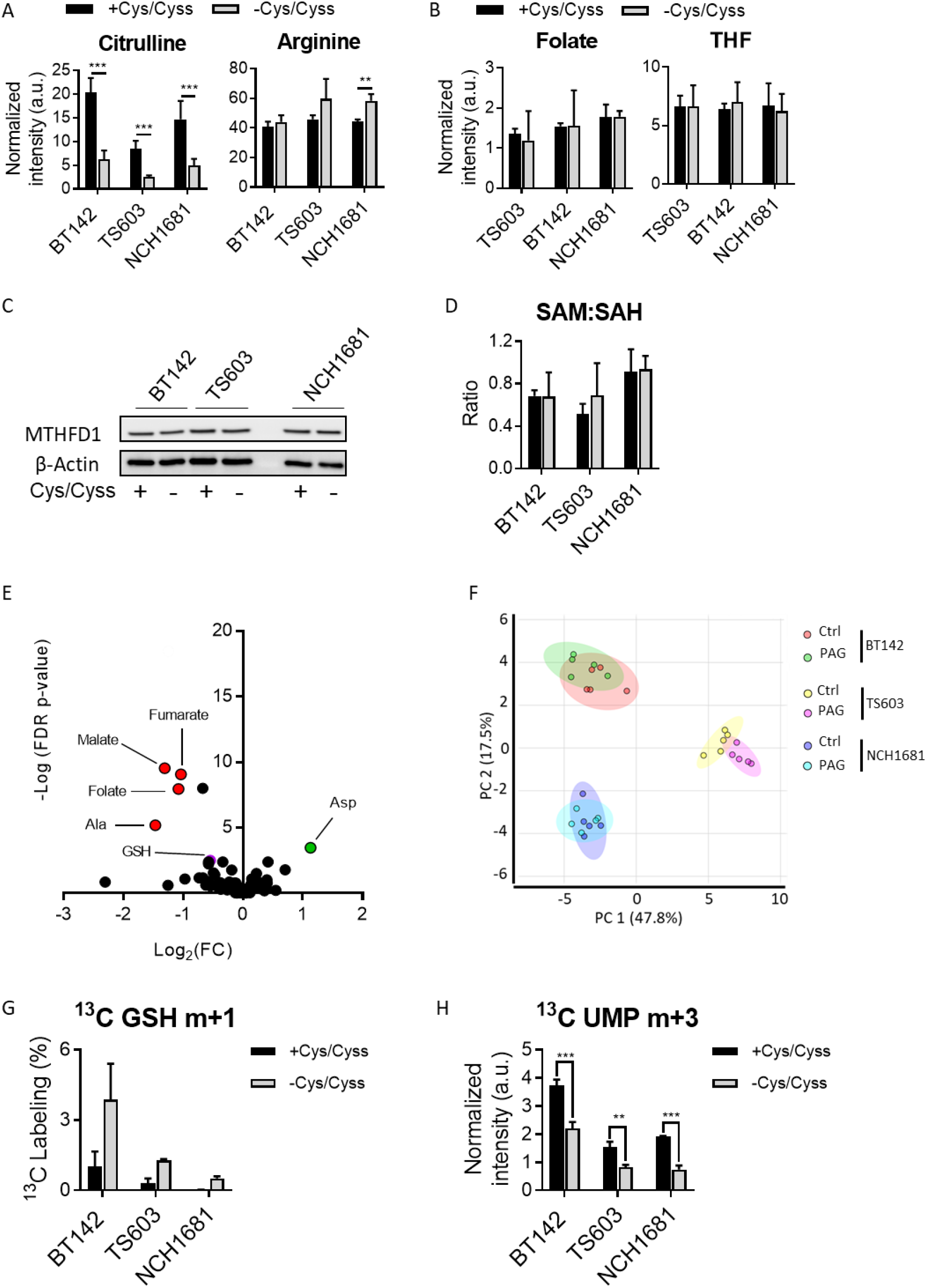
(A) Levels of citrulline, arginine, and (B) tetrahydrofolate (THF) and folate for the 3 cell lines grown in full and cysteine/cystine-lacking media for 48 hours. Metabolite levels are computed from the LC-MS metabolic profiling experiment (n = 5, bar plots show mean values ± SD, *p* values from a *t*-test adjusted for multiple comparisons by the FDR method. \*\**p* < 0.005; \*\*\**p* < 0.001). (C) Western blot of methyltetrahydrofolate dehydrogenase 1 (MTHFD1) as a marker of folate cycle activity. (D) Ratio of S-adenosylmethionine (SAM) and S-adenosylhomocysteine (SAH) for the 3 IDH1-mutant glioma cell lines computed from the LC-MS metabolic profiling experiment (n= 5, bar plots show mean values ± SD). (E) Volcano plot of the glioma cell lines treated with 1 mM propargylglycine (PAG) vs. controls. Only metabolites that surpassed the thresholds established as FDR-corrected *p* < 0.05 and Log_2_(FC) > 1 or < −1 are highlighted (red, downregulated metabolites; green, upregulated metabolites) in addition to glutathione (GSH). (F) PCA scores plot using Log-transformed metabolites levels detected by LC-MS; 95% confidence ellipses are shown as colored ovals for each cell line and condition (treatment with 1 mM PAG). (G) Percentage of m+1 glutathione (GSH) isotopologue over the total pool of GSH from ^13^C-C3-serine (bar plots are mean ± SD, n = 3). (H) ^13^C incorporation from ^13^C-U-glutamine into uridine monophosphate (UMP) as the m+3 isotopologue (bar plots are mean ± SD, n = 3; \*\**p* < 0.005, \*\*\**p* < 0.001, two-tailed Student’s *t*-test).

**Figure S3:**
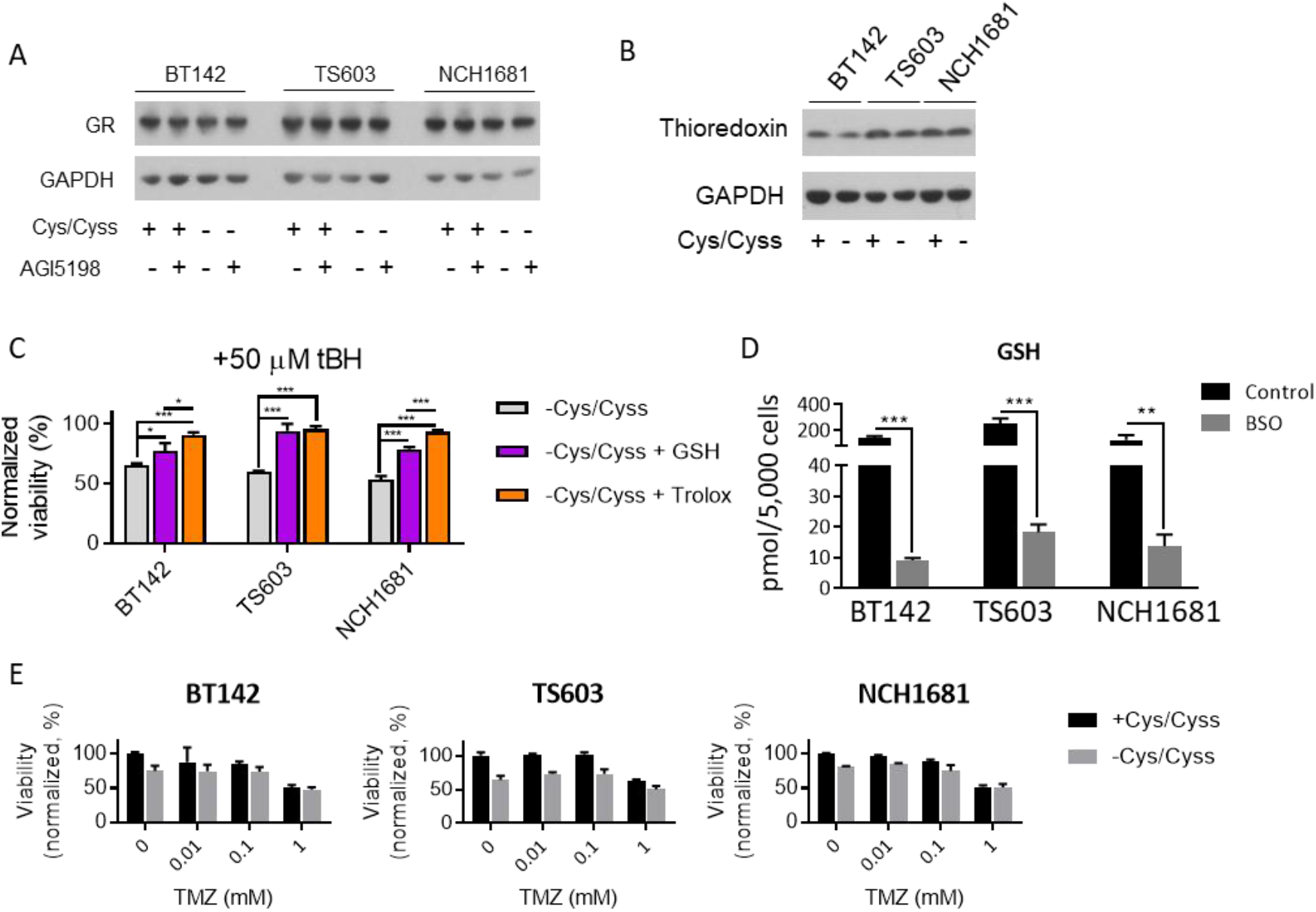
(A) Western blot of glutathione reductase at the different conditions depicted in the figure. (B) and thioredoxin. (C) Normalized viability of the 3 IDH1-mutant glioma cell lines treated with 50 µM of tBH for 24 hours in medium spiked with 0.1 mM of GSH or 0.25 mM Trolox (bar plots are mean ± SD, n = 3; \**p* < 0.05, \*\*\**p* < 0.001, one-way ANOVA followed by Tukey’s multiple comparisons test). (D) Quantification of GSH levels after treatment with 250 µM BSO (bar plots are mean ± SD, n = 3; \*\**p* < 0.005; \*\*\**p* < 0.001, two-tailed Student’s *t*-test). (E) Normalized viability of the 3 IDH1-mutant glioma cell lines treated with temozolomide (TMZ) for 96 hours in full medium and medium lacking cysteine/cystine. No significant differences (by two-way ANOVA) were found between the cells grown in medium lacking cysteine/cystine and the combination with any of the TMZ concentrations.

**Supplementary Figure 4:**
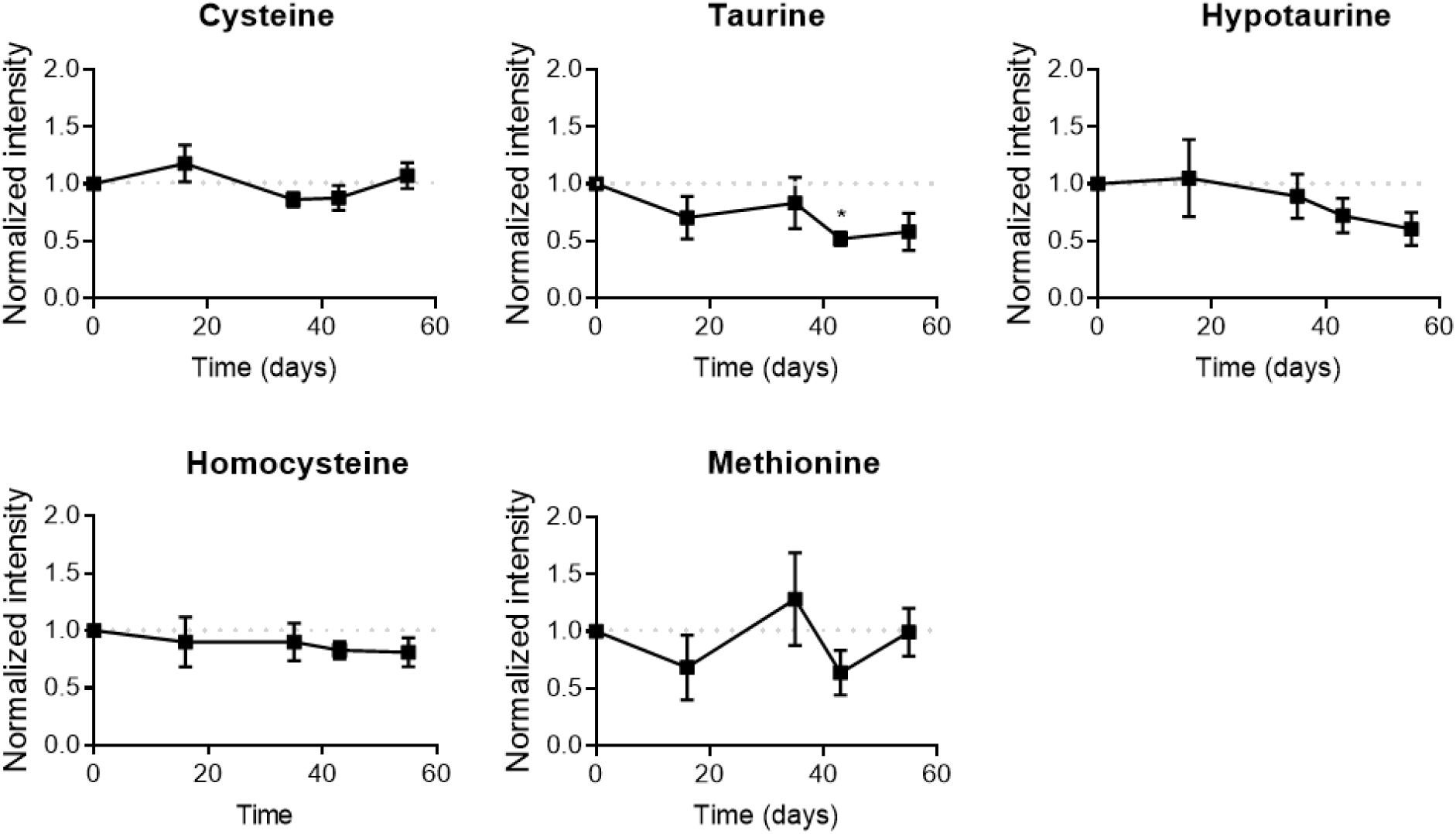
Normalized intensities of cysteine-related metabolites in plasma collected from a mouse model of glioma and normalized to those computed for the control group (dotted line, reference values of the control group; metabolite levels for the cysteine/cystine-free diet group are shown as mean ± SEM; n = 5; \**p* < 0.05 by two-tailed Student’s *t*-test with Welch’s correction).

